# Physiological consequences of acute heat exposure in mid-gestation on placental, foetal and maternal blood flow using a mouse model

**DOI:** 10.64898/2026.04.06.713526

**Authors:** Shannon C. Francis, Colin E. Murdoch

## Abstract

Pregnant women are particularly susceptible to adverse outcomes from environmental heat, yet the physiological effects of acute heat exposure during pregnancy remain poorly understood. Some physiological changes are monitored in humans; however, investigation of underlying molecular mechanisms requires invasive methods that can only be ethically applied in mammalian models. Moreover, research with animal models has largely focused on early and lethal teratogenic effects of heat exposure and lacks longitudinal physiological monitoring, detailed parameterisation of heating regimes and in-depth investigation of underlying mechanisms.

Here we used a mouse model to investigate the impact of a controlled acute heat exposure at mid-gestation (E12·5), slowly elevating core body temperature (CBT) over 210mins to raise CBT by ∼1°C. Using high-frequency ultrasound and morphological analyses, we observed delayed alterations in placental and foetal cerebral blood flow indicative of a brain-sparing response, alongside reduced placental labyrinth zone size. Additionally, maternal cardiac function was impaired, accompanied by cardiac and renal fibrosis and elevated circulating soluble Flt-1 levels, an anti-angiogenic biomarker of gestational hypertension. These findings demonstrate that brief heat stress at mid-gestation can induce lasting effects on placental function and maternal cardiovascular health in a mammalian model, highlighting potential risks for pregnancy outcomes under increasing global temperatures.

Together this data suggests that an acute exposure to heat elevating core body temperature by 1·2°C can induce a long-term impact on both placenta and maternal health in a mouse model. It will be important to understand the molecular changes which underpin the pathophysiology and whether this is translated to humans.

## Introduction

As the world warms, there is an urgent need to determine the impact of heat-stress on pregnancy outcomes ^1–4^. Pregnant women are one of the most vulnerable groups that are likely to be most impacted by global climate change, with clear evidence from across the globe that environmental heat correlate with enhanced risk of still or preterm birth and growth-restriction, despite our homeothermic nature ^5–7^. Similarly, heat waves cause an increase in admissions for both maternal and foetal complications^8^. Physiological and anatomical adaptations hinder the ability to adequately thermoregulate during pregnancy, as a result of significant increases in cardiac output, body fat, and metabolism^9^. This combination of haemodynamic changes, increased metabolic load and a decreased body mass to surface area ratio make it difficult to dissipate heat, thus increasing maternal core body temperature^10^. Approximately 85% of foetal heat is transferred across the placenta, driven by a temperature gradient where foetal temperature is ∼0·5°C higher than maternal CBT^11^, consequently increasing the risk of teratogenic temperatures for the embryo in response to maternal heat exposure. Nonetheless, the consequences of extreme heat exposure in pregnancy are poorly understood.

Exposure to heat in the first trimester also correlates with morphological abnormalities, including neural, facial, and cardiovascular defects, which align with known critical periods in tissue morphogenesis^12–14^. Mammalian models using varied heating regimes and analytical methods, and focussing largely on organogenesis stage embryos, confirm these tissue vulnerabilities following even brief exposure (>2·0°C normal body-temperature)^15^. The impact of heat on the placenta has been postulated to be secondary to peripheral vasodilation, which shifts blood flow away from the placenta and foetus ^4,16^. Indeed, chronic heat exposures in varied animal models display reduction in uterine and umbilical flow ^17,18^, enhanced vascular resistance^19^, disrupted placental barrier function and vascularisation^11,19^. Such changes are hallmarks of placenta dysfunction in preeclampsia and foetal growth restriction. Overall, these findings support the hypothesis that extreme heat triggers placental dysfunction leading to changes in placenta vascularisation, and consequent ischemia/hypoxia and cellular stress pathway induction in both the placenta and embryo, aligning with mechanisms that underpin the Developmental Origin of Health and Disease (DOHaD) theory^20^. How heat-induced placental dysfunction impacts development of the placenta and embryo and how this differs due to heat duration, developmental stage, and following prior maternal heat-acclimation remain to be determined.

Here, we aimed to establish a mouse model of acute heat exposure (AHE) mid-gestation, to begin to explore the pathophysiological and mechanistic impacts of heat exposure during pregnancy on maternal and foetal health.

## Methods (including Ethics, Statistics, and Role of Funders)

### Ethics statement

All animal procedures were carried out in accordance with UK Home Office Guidance on the Operation of the Animals (Scientific Procedures) Act 1986, with approval by the University of Dundee Welfare Ethics committee under project license PP7330886.

### Housing and Timed pregnancy

CB57BL/6j mice (12-16 weeks (Charles River)) were group housed (3-5 mice per cage) at a temperature of 18-22°C and humidity 45-65% with a 12-hour light/dark cycle. Dams were timed mated in singles or pairs with male studs for 18-hours. Dams were checked daily for vaginal plugs to confirm pregnancy along with regular weighing. The presence of a vaginal plug signified gestation day E0·5.

### Randomisation and Blinding

Upon confirmation of pregnancy, dams were randomly assigned to either “Heat” or “Control” groups in an alternating manner to maintain even distribution between experimental groups. Each dam was given a unique sample number, and all data acquisition and analyses were completed blinded. Except for core body temperature measurements where the operator knew if the mice were in the heat of control group.

### Acute heat exposure heating regime

At gestation E12·5, mice in the “Heat” group were singly housed and underwent an acute heating regime. Singly housed mice were placed in a pre-warmed (37·5°C) heating chamber (Scanbur). Mice were housed singly for the heating regime, with gradual increases in temperature from 37·5°C to 40·5°C over a three-hour period (Fig.1B). CBT was recorded via rectal probe (VisualSonics) every 60 minutes. Control mice underwent the similar procedure except they were not placed in the heating chamber.

**Figure. 1.**
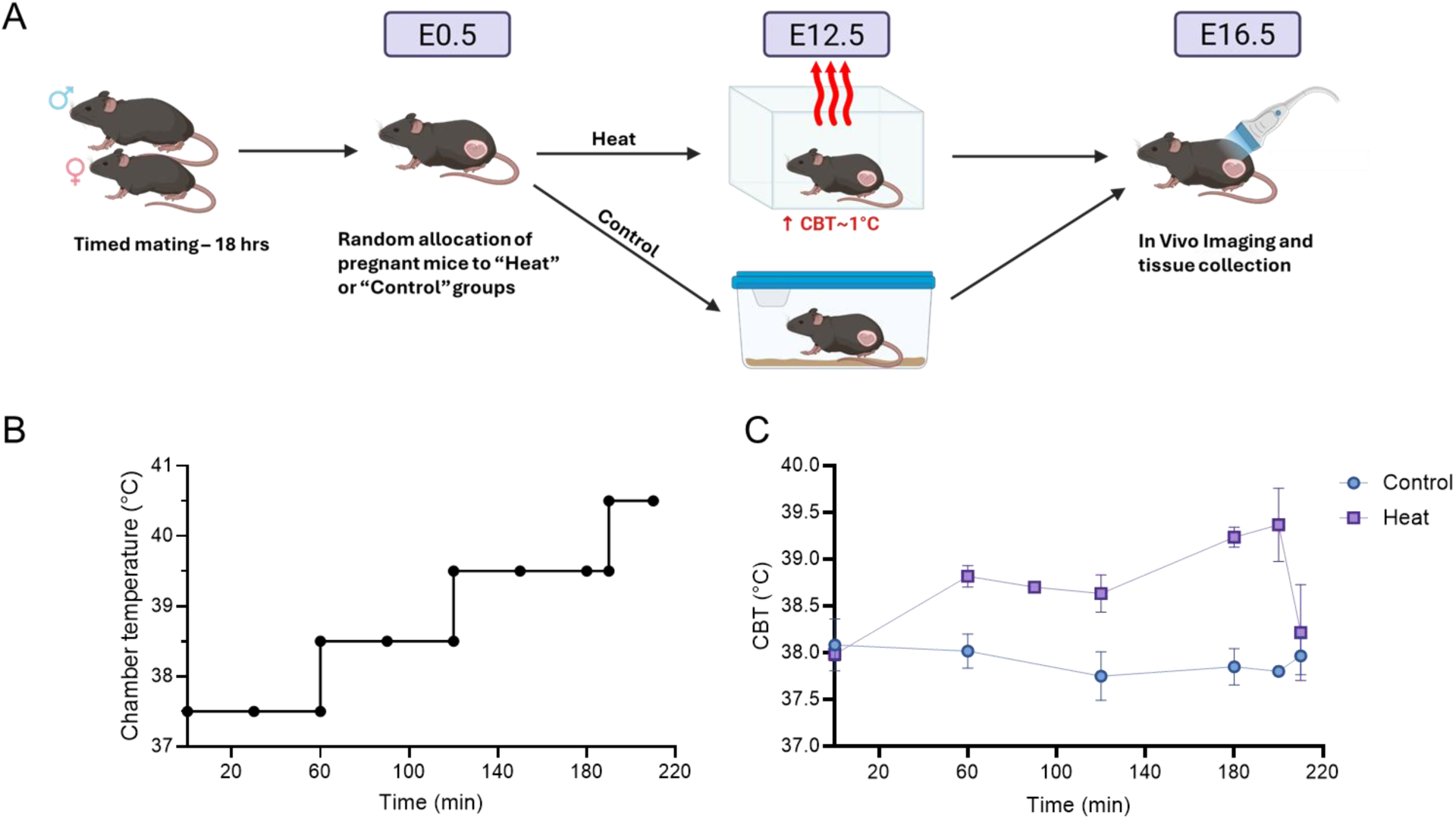
Acute Heat Exposure (AHE regime induced an increase in core body temperature (CBT). (**A**) Schematic of experimental procedure, with time-mated mice being randomly allocated heat group at gestation E12·5. Ultrasound combined with pulse wave Doppler and subsequent harvesting at E16·5. **(B)** Heating chamber was initially set at 37·5°C for the first 60 minutes and was increased by 1°C every 30-60 minutes until the final increase to 40·5C at 190 minutes for 10 mins. **(C)** CBT was measured regularly during the heating procedure by digital rectal probe (*n*=12).

### Pulse Wave Doppler analysis

Ultrasound coupled with Pulse wave Doppler (VevoF2, VisualSonics, Toronto, CA) analysis was conducted on all mice at 4-days post-heat or control. Dams were anaesthetised (Isoflurane, 1·5-2%) and placed in a supine position on a heated platform. The electrocardiogram monitored heart and respiratory rate, and a rectal probe measured core body temperature throughout the procedure.

The uterine artery (UtA) was identified using the bladder as a landmark. Each conceptus was identified in relation to the bladder, and the umbilical artery (UmA) Doppler measurements were obtained proximal to the placenta. Doppler measurements of foetal middle cerebral artery (MCA) were obtained using a cross-sectional view of the head. Foetal and placental blood flow profiles were analysed for the conceptus closest to the bladder.

Pulse wave Doppler analysis was carried out in VevoLab F2 software. Measurements were averaged across three consecutive waveforms per image and used to calculate the cerebroplacental ratio (CPR) and uterine PSV ratio by the following formulae:

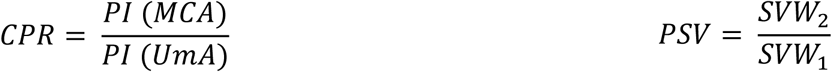

### Echocardiography

Left-ventricle long-axis echocardiography was performed at E16·5 to assess maternal cardiac function. The endocardium was traced from the left ventricular outflow tract to the apex during systole and diastole to calculate cardiac output, ejection fraction, fractional shortening, end diastolic volume, and end systolic volume. Interventricular septum thickness (IVS) was measured at end diastole and systole from M-Mode images. Measurements were averaged across three cardiac cycles.

### Tissue harvest and measurements

After ultrasound analysis at E16·5, cardiopuncture was performed under anaesthesia (Isoflurane 1·5-2%). Tissues from the dams were collected and embryos and placentas were dissected from their amniotic sacs. All embryos, placentas and maternal organs were weighed and fixed or snap frozen. Foetal morphometrics were recorded post-fixation using callipers.

### Sex genotyping

Sex genotyping was carried out on DNA extracted from embryonic tail samples^21^. Briefly, multiplex polymerase chain reaction (PCR) was conducted (GoTaq® MasterMix, annealing temperature 58°C) using two sets of primers, targeting the sex determining region Y (*Sry*) gene (Sequence 8276–8295 5′-TGGGACTGGTGACAATTGTC-3′ and 8677–8658 5′-GAGTACAGGTGTGCAGCTCT-3′) and *IL3* gene (Sequence 792–801 5′-GGGACTCCAAGCTTCAATCA-3′ and 1335–1316 5′-TGGAGGAGGAAGAAAAGCAA-3′) respectively. Samples were loaded on a 3% agarose gel and where female samples displayed a single band (544bp), and male samples showed two bands (402bp, 544bp).

### Histology

Dam organs and placentas were immersion-fixed in 4% paraformaldehyde and processed into paraffin wax. Samples were sectioned to 6µm on a Shandon microtome (Thermo Fisher Scientific) and mounted on Superfrost Plus microscope slides.

Placental sections were deparaffinised and rehydrated and washed under running water for 3 mins. Slides were stained with diluted Mayer’s haematoxylin (2:1 dilution, 3 mins). After washing, slides were incubated with Scott’s tap water (2% magnesium sulphate, 0·25% sodium bicarbonate, 10 seconds) and counterstained with eosin (3-5 seconds). Slides were rinsed, dehydrated and cleared and mounted. Image acquisition was performed using Invitrogen EVOS M7000 Imaging System. Images were taken at 4x magnification and automatically stitched together to visualise entire tissue section.

Quantitative analysis was carried out to assess changes in placental architecture. Samples that were not in a comparable orientation were excluded from analysis. Images were analysed blindly in QuPath software where the boundaries of each placental zone were determined manually by changes in morphology. Each zone was measured and converted into percentage areas.

To determine cardiac and renal collagen deposition, sections of maternal left ventricle and kidney were stained with picrosirius red. Slides were deparaffinised, rehydrated and soaked in ddH_2_O (5 mins). Slides were stained in PicroSirus Red (PcSR) (0·5g Direct Red in 500ml picric acid, 1 hour) and washed in 0·5% acidified water (0·5% acetic acid, 2 mins) before being dehydrated, cleared and mounted. Slides were imaged using Invitrogen EVOS M7000 Imaging System at 10x magnification.

Quantitative analysis was carried out blindly in ImageJ to assess the level of kidney and left ventricle fibrosis. All images were calibrated using the scale bar which was subsequently removed to avoid being included in the analysis. Fibrosis was quantified using images at 10x magnification and splitting the image into RGB stacks. The green channel was used, and the threshold was adjusted to match the areas of fibrosis. Multiple images were analysed per sample and overall average percentage area of PcSR collagen staining.

### Enzyme-Linked Immunosorbent Assay (ELISA)

ELISA was performed on maternal serum to measure circulating levels of sFlt-1 according to the manufacturer’s instructions (R&D Systems Mouse sVEGF R1/Flt-1 DuoSet ELISA, Cat #DY471). A standard curve was generated using linear regression by plotting the mean absorbance of the standards against the log_10_ of the concentration. Using the absorbance of the unknown sample data, their concentration was interpolated from the standard curve.

### Statistical analysis

Statistical analyses were conducted in GraphPad Prism version 10·4·2. Data is denoted as mean ± standard error of the mean (SEM) unless stated otherwise. All data was assessed for normality before conducting either Mann-Whitney U tests or Student-T tests for non-parametric and parametric tests respectively. Statistical significance was set as P<0·05.

## Results

### Acute extreme heat exposure

Pregnant, timed mated C57BL6j mice were randomly assigned to either heat or non-heat control groups (Fig.1A). At E12·5, mice in the heat group underwent acute heat exposure (AHE) by placing into an open cage inside a prewarmed heating chamber (37·5°C). Rectal core body temperature (CBT) was monitored regularly as the heating chamber temperature was gradually increased from 38-40·5°C over a period of 3 hours (Fig.1B) in order to raise CBT by ∼1.2°C. Control (non-heated) mice (control) also had CBT monitored but were not placed in the heating chamber. The baseline CBT of pregnant mice (E12·5) was found to be 38+/-0·14°C (*n*=6) (Fig.1C). During the first hour of heating there was an initial increase in CBT of approximately 0·5°C. A further increase in CBT up to 1·2°C was achieved after heating for an additional 140 mins (Fig.1C). All mice returned to their pre-heat CBT as soon as they were removed to room temperature (21°C) (Fig.1C).

### AHE impact on foetal and placental blood flow inducing a potential ‘brain sparing’ reflex

We first evaluated whether AHE induced changes in blood flow within the maternal (Fig.2A), placenta (Fig.2B) and foetal (Fig.2C) vascular beds, using high frequency ultrasound (VisualSonics) and pulse wave Doppler. The pulsatility index (PI) which reflects the resistance to blood flow was recorded 4-days post-heat. When assessing maternal circulation, the PI in uterine artery (**UtA**) (maternal) (P=0·73) and maternal heart rate (HR) (P=0·21) were similar between control and heat groups (Fig 2D-G). There was no significant difference in other pulse wave Doppler parameters, including the PSV ratio (Fig 2J). To assess the impact of AHE on placenta and foetal blood flow we measured umbilical artery (**UmA**) PI. AHE induced a significant increase in UmA PI 4-days post-heat (P=0·03) (Fig 2E), suggesting an increase in resistance in blood flow from the foetal side of the placenta. In contrast, there was a decrease in PI in foetal middle cerebral artery (MCA) post-heat (P=0·02) compared to control (Fig 2F). The changes in UmA and MCA PI did not alter the foetal heart rate (Fig 2H-I), providing confidence in the pulse wave Doppler measurements. The cerebroplacental PI ratio (**CPR**) is used clinically to assess placenta dysfunction and is indicative of brain sparing reflex. The CPR was significantly lower in the heat group (0·8+/- 0·10) compared to control (1·0+/- 0·18, Fig.2K) 4-days after AHE.

**Figure. 2:**
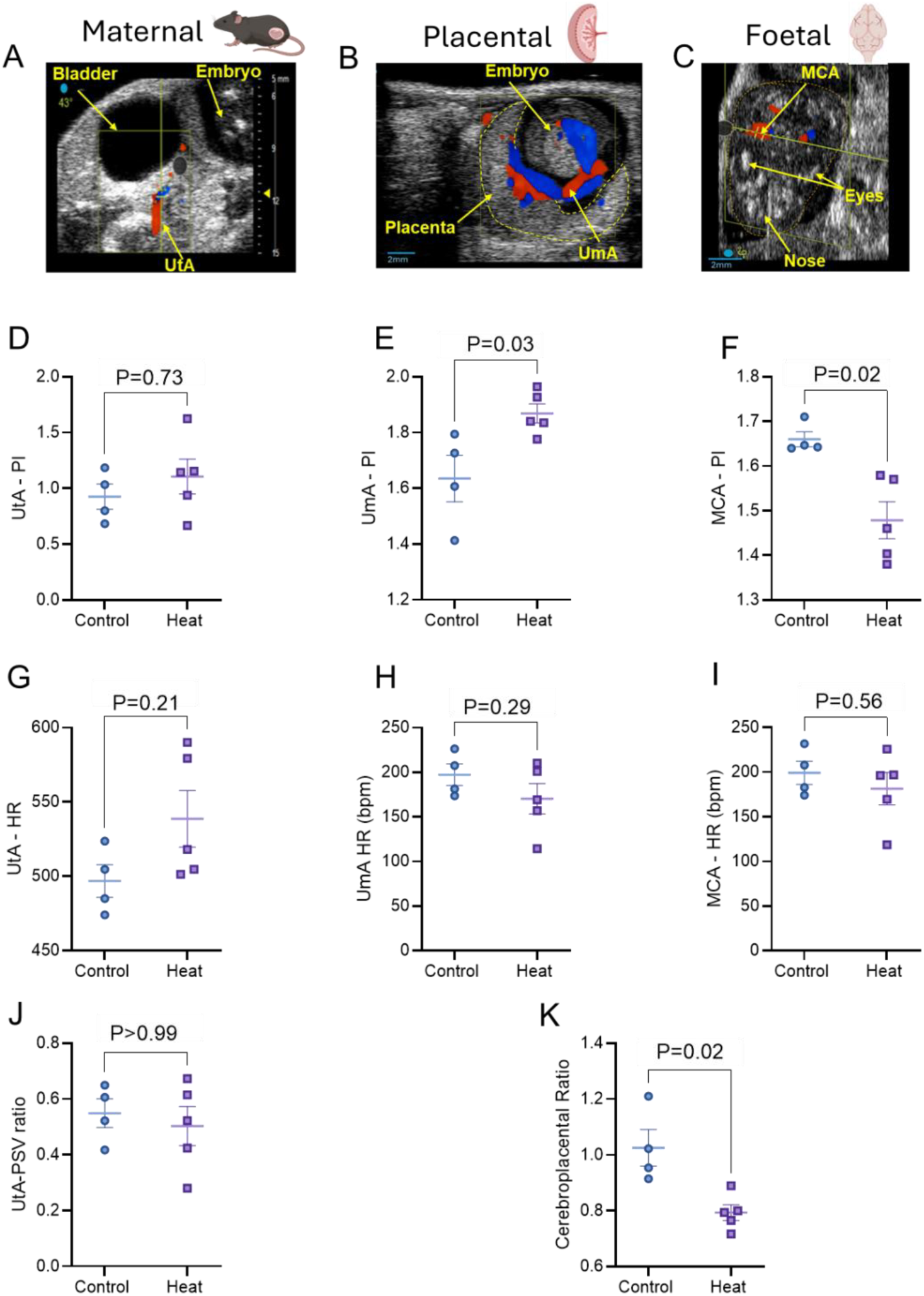
Impact of Acute Heat Exposure (AHE) on maternal, placental and foetal vascular beds. High frequency ultrasound and colour doppler images used to identify the blood flow in maternal (A) uterine artery (UtA), (B) umbilical artery (UmA) and (C) foetal middle cerebral artery (MCA). Blue and Red colour Doppler relates to the respective away or toward direction of blood flow in relation to the ultrasound probe which is positioned at the top of each image. Pulsatility index (PI) from pulse-wave Doppler blood flow profiles in UtA **(D)**, UmA **(E)** and MCA **(F)** in control (*n*=4) and heat (*n*=5). Corresponding heart rate (HR) n UtA **(G),** UmA **(H)** and MCA **(I)**. UtA peak systolic velocity ratio (**J**) index was calculated as a ratio of the two systolic velocity waves. Cerebroplacental ratio index **(K)** calculated as a ratio from MCA and UmA PI. Non-parametric Mann-Whitney U tests were conducted for all comparisons and data presented as mean ± SEM. Statistical significance was set at P<0·05. *N*=4 “Control” and *n* =5 “Heat” where each *n* represents one mouse.

### The consequence of AHE on pregnancy outcome

To assess the impact of AHE on pregnancy outcome, all pregnancies were terminated four days post-AHE (E16·5). There were no significant changes in reabsorption rates between control and heat groups (P=0·39) (Fig.3F). All pups underwent sex genotyping for the presence of male-specific gene Sry. There was equal male-to-female ratio in the pups from each litter (Fig.3G). Foetal and placental weights were recorded, and the respective ratio was calculated as a measure of placental sufficiency, which did not change between the groups (Fig.3H). In contrast, AHE induced changes in placenta architecture, observed by a 10·7% reduction in labyrinth zone area (P<0·0001) (Fig.3C) and a 10·3% increase in junctional zone area (P=0·01) (Fig. 3D). There was no significant change in decidua zone area (P=0·40) (Fig.3E).

**Figure. 3:**
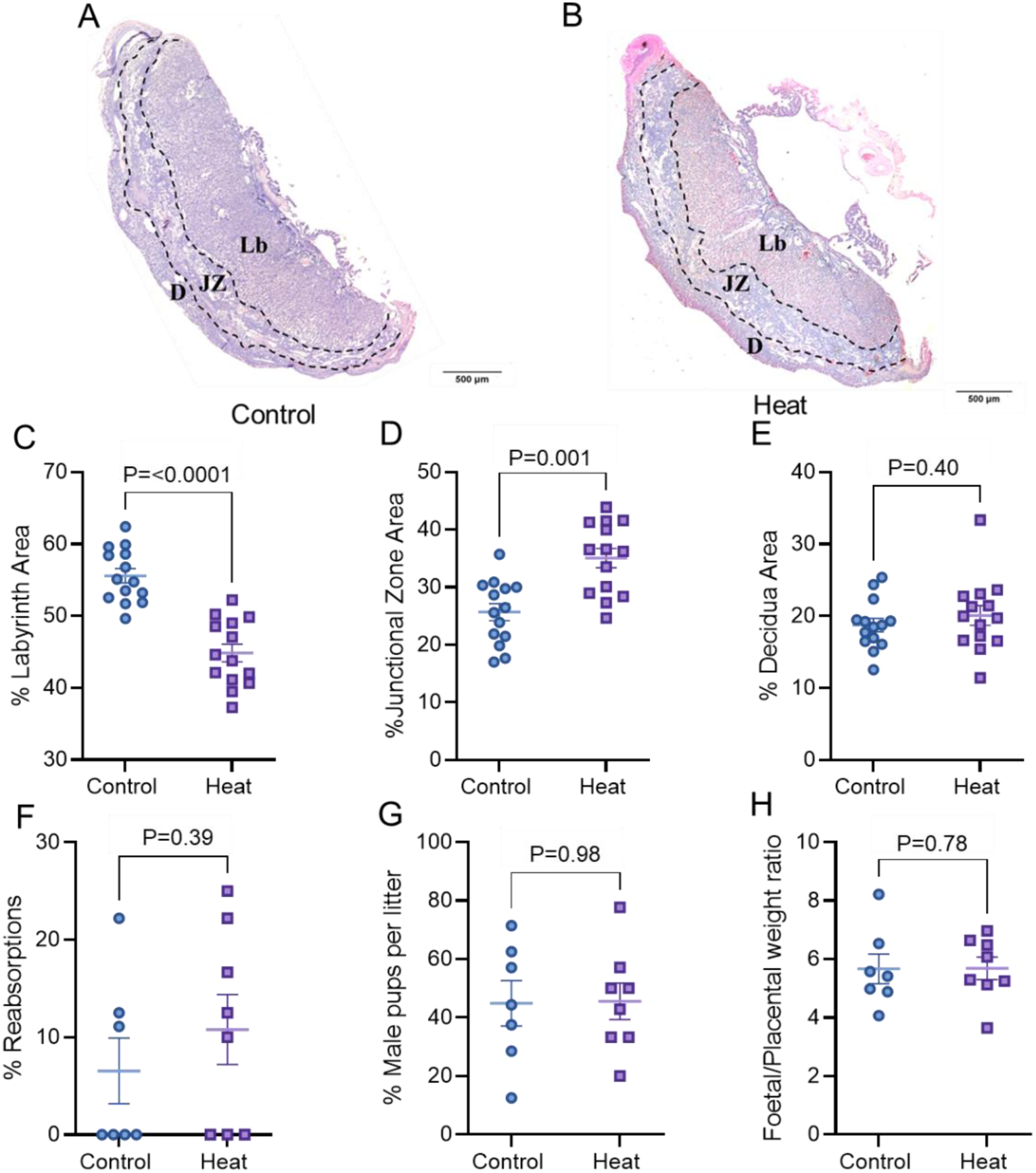
Comparison of foetal morphometrics and pregnancy outcomes between heat and control groups at E16·5. Representative H&E-stained placenta sections from Control **(A)** and Heat **(B)** groups, 4x magnification, scale bar 500μm. Differences in percentage area of labyrinth **(C)**, junctional (**D**) and decidua **(E)** zones were evaluated. Non-parametric Mann-Whitney U tests were performed with statistical significance set as P<0·05. All data presented as mean ± SEM, data points represent individual placentas where *n*=7 “Control” and *n*=8 “Heat”. *Abbreviations: D = decidua, JZ = junctional zone, Lb= labyrinth.* % Reabsorption in control and heat litter to assess embryonic loss in control (*n*=7) and heat (*n*=8) groups at E16·5 **(F)**. % of male pups per litter in control (*n*=7) and heat (*n*=8) groups **(G).** Average foetal (C) and placenta weight (D) control groups. Non-parametric Mann-Whitney U were performed with statistical significance set as P<0·05. All data presented as mean ± SEM, each data point represents average measurements across a single litter.

### AHE induces maternal cardiac and renal dysfunction

In addition to assessing foetal and placenta outcomes, we also investigated the impact of AHE on maternal cardiovascular health. Left-ventricle long-axis echocardiography was performed at E16·5, four days after AHE, revealing a clear decrease in cardiac function. The heat group showed decreases in fractional shortening (Fig.4D), ejection fraction (Fig.4C), and cardiac output (Fig.4B). Importantly, the heart rate during these measurements was similar between control and heat groups (Fig.4K). There were no differences in end diastolic volume (Fig.4G), end systolic volume (Fig.4F), stroke volume (Fig.4E), or cardiac dimensions (intraventricular septal thickness (Fig.4I-J).

**Figure. 4:**
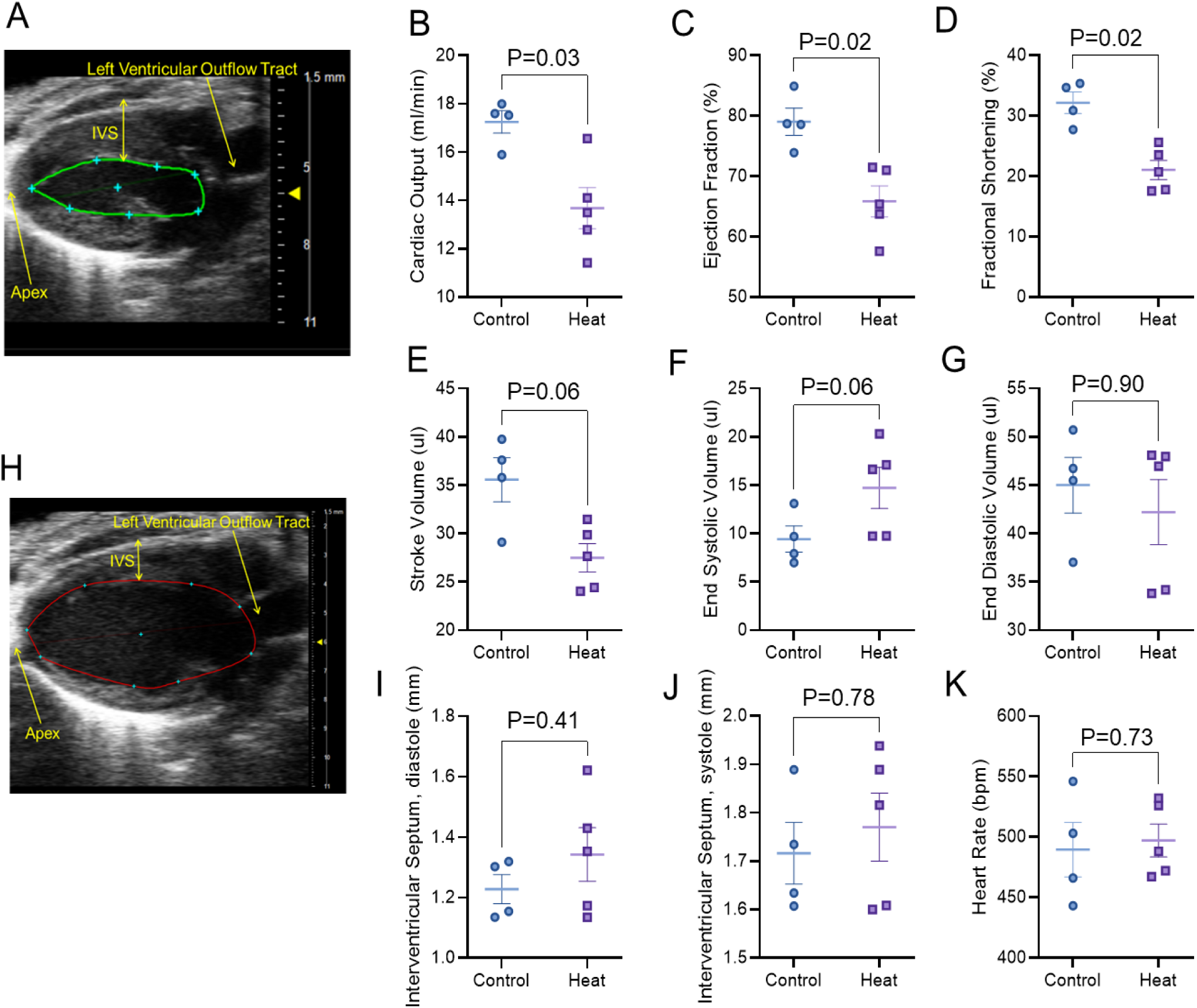
AHE-induced maternal cardiac dysfunction. Long axis echocardiography images taken during systole **(A)** and diastole **(H)**. Echocardiographic assessment of maternal left ventricle function in control (*n*=4) and heat (*n*=5) mice for cardiac output **(B)**, ejection fraction **(C)**, fractional shortening **(D)**, stroke volume **(E)**, end systolic volume **(F)**, end diastolic volume **(G)** and cardiac heart rate **(K)**. Intraventricular septal thickness obtained from m-mode LV imaging in anesthetized mice (Isoflurane 1·5-2%) during full diastole **(I)** and systole **(J)**. Non-parametric Mann-Whitney U were performed with statistical significance set as P<0·05.

Maternal organs were weighed immediately after excision and normalised to tibia lengths to account for variations in body weight between mice. Mice that underwent heat at E12·5 had significantly larger hearts (Fig. 5A), with no significant differences in the weight of kidney, lung or liver (Fig.5B-D respectively). We next investigated maternal cardiac and renal fibrosis by staining collagen with Sirus Red. In both heart (Fig.5G) and kidney (Fig. 5J) there was a significant increase in Sirus Red collagen staining in the heated group compared to controls. The heat group compared also had higher plasma levels of sFLT-1, an antiangiogenic factor (sFLT-1) (Fig.5K) used clinically as a blood biomarker of gestational hypertension and thought to be released from dysfunctional placenta (^22,23^).

**Figure. 5:**
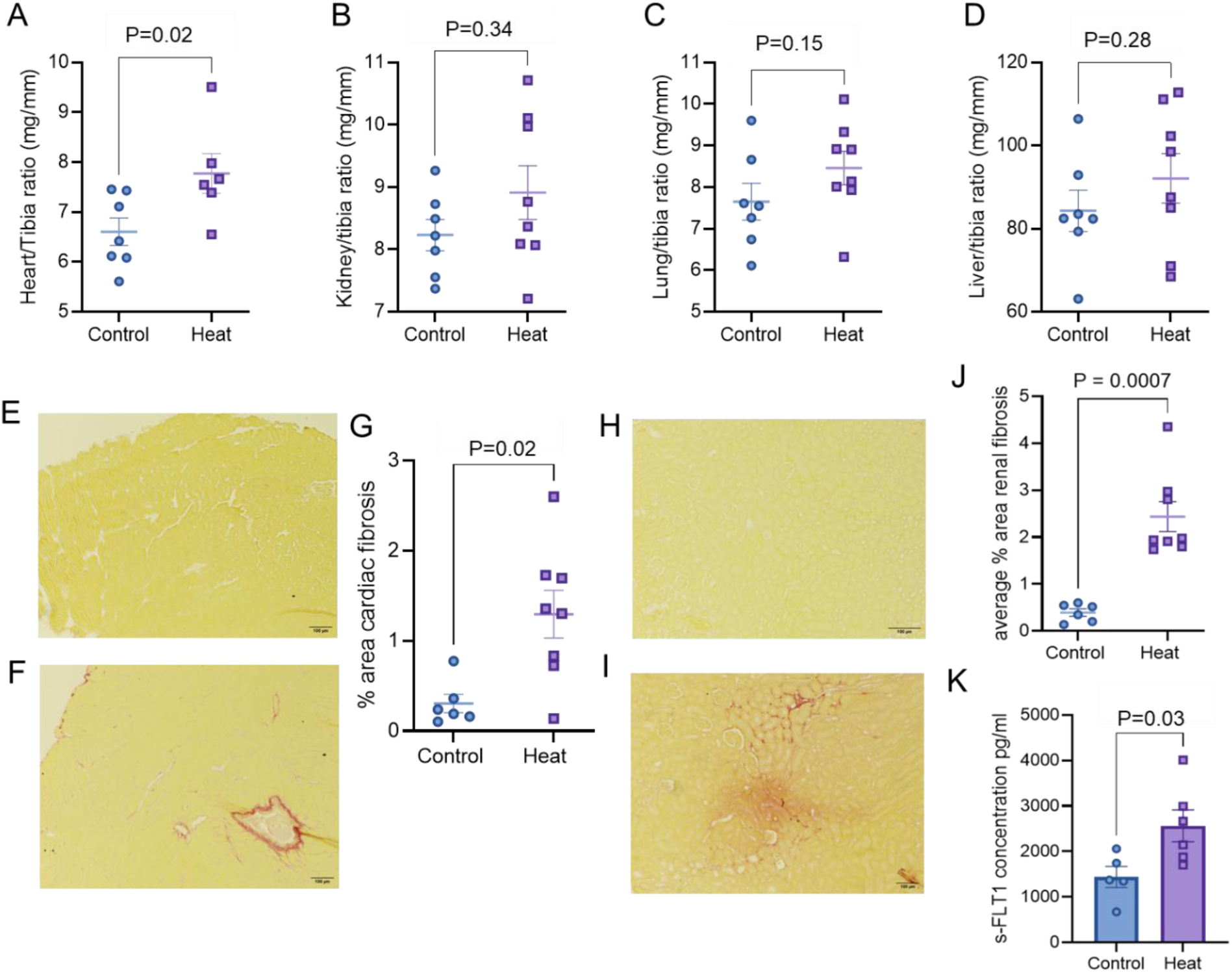
Assessment of cardiac fibrosis in heat-exposed mice during pregnancy. Organ weights were normalised to tibia measurements, presented as ratios, to account for differences in body size maternal heart **(A)**, kidneys **(B)**, lungs **(C)** and liver **(D)** were recorded after excision Representative picrosirius red staining of maternal left ventricle **(E&F)** and kidney **(H&I)** in control **(E&H)** and acute heat-exposed **(F&I)** mice. Percentage area of cardiac **(G)** and renal **(J)** fibrosis. All data presented as mean ± SEM where each data point represents one mouse. Mann-Whitney U were performed with statistical significance set as P<0·05. All data presented as mean ± SEM, *n*=12 “Control”, *n*=9 “Heat” where each n is one mouse. Mice who underwent the acute heating regime exhibited higher levels of sFlt-1 in maternal plasma compared with control mice **(K)**. Non-parametric Mann-Whitney U were performed with statistical significance set as P<0·05. All data presented as mean ± SEM, *n*=6 “Control”, *n*=5 “Heat”.

## Discussion

We have established a mouse model of acute heat exposure with the ability to track *in vivo* physiological changes which opens up the potential to correlate these with placental molecular signatures.

Clinically, ultrasound imaging of uterine, umbilical, and foetal vasculature in human pregnancies is instrumental in the prognosis of pregnancy complications and adverse foetal outcomes. Placental insufficiency is normally characterised by increased blood flow resistance through the uterine artery thus limiting the availability of oxygen and nutrients to the foetus^24,25^, however our data showed no changes in uterine artery Doppler parameters. Despite this, rodent models of IUGR by hypoxia ^26,27^ and cohort studies of pregnancy outcomes during heat stress^28^ have reported similar findings, suggesting this could indicate a compensatory downstream vasodilatory response to preserve sufficient placental perfusion. Nevertheless the maintenance of pulse wave doppler blood flow in the uterine artery demonstrates a preserved maternal vascular response. In contrast, AHE significantly impacted both umbilical and middle cerebral artery pulsatility index, contributing to a decreased cerebroplacental ratio in heat-exposed mice. Reduced PI and therefore CPR indicates decreased blood flow resistance to the foetal brain, which is suggestive of the adaptive “brain sparing” response. This is often observed in cases of foetal hypoxia or IUGR, inducing a haemodynamic redistribution of blood flow towards the brain to conserve neurodevelopment^29^. Although brain sparing is often recognised as a protective response, it cannot fully mitigate prolonged placenta insufficiency. The redistribution of cardiac output away from the peripheral organ systems has been shown to result in adverse foetal outcomes, impaired neurodevelopment and increased predisposition to chronic illnesses in both preclinical and cohort studies^30–32^.

In the current study, placentas from heat-exposed mice exhibited altered placental architecture, evidenced by a decrease in the labyrinth zone and increased junctional zone. The labyrinth zone is the site of maternal-foetal exchange within the placenta and continues to increase in size throughout gestation to support the growing demands of the foetus. Our results suggest that this normal growth is inhibited under heat stress, resulting in a smaller surface area for maternal-foetal transport and exchange, as well as further disrupted angiogenesis by the upregulation of sFlt-1. Similar results along with reduced blood spaces have been observed in animal models of preeclampsia and IUGR, indicative of hindered nutrient transport to the foetus^33,34^. Despite significant changes in the maternal-foetal interface, there were no differences in the number of reabsorbed embryos or placental sufficiency between the groups. As foetal growth largely happens towards the end of pregnancy, it is likely that altered foetal and placental morphometrics would not be observed at E16·5. Future studies should aim to investigate foetal outcomes as a later gestation.

Pregnancy requires extensive cardiometabolic adaptation to facilitate proper foetal growth and development. The cardiovascular system is the first organ system to be affected by heat stress^35^ which is likely exacerbated during pregnancy. Here, we found that AHE greatly impacted overall cardiac function, shown by echocardiographic assessment of the maternal left ventricle. Reductions in cardiac output, fractional shortening, and ejection fraction indicate impaired systolic function and contractility, which could contribute to altered uteroplacental blood flow. Similarly, heated mice presented with increased heart weights in comparison to control mice, whilst there were no changes in other organ weights. Physiological hypertrophy is expected during pregnancy as a haemodynamic adaptation to facilitate proper foetal growth and development; however, these results suggest that there is considerable cardiac strain in response to heat exposure. In our model, we also observed increased cardiac and renal fibrosis in heat-exposed dams which were predominantly perivascular, consistent with our findings of compromised cardiovascular function and increased cardiac demand. Increased collagen deposition within the kidney is often indicative of acute kidney injury, a common consequence of hypertensive disorders of pregnancy. Animal models of induced preeclampsia have observed glomerular endotheliosis, increased collagen deposition, proteinuria, and an overall decline in renal health^36,37^ which is consistent with our findings and the indication of a preeclampsia-like phenotype.

As anticipated, there was a significant increase in the plasma levels of sFlt-1 in the maternal circulation of heated dams compared with the control group. The sFlt-1/PlGF ratio has been well-reported as a clinically used metric for assessing preeclamptic risk, signifying the balance between pro- and anti-angiogenic factors. This rise in circulating sFlt-1 supports the hypothesis that heat stress disrupts normal angiogenesis, leading to endothelial dysfunction, oxidative stress and affecting overall pregnancy outcomes.

This study is unique in that we were able to align physiological differences in blood flow and vascular metrics to morphological changes in individual tissues. We believe our mouse model has physiological relevance to humans as the mouse model is well-characterised for pregnancy studies which allows us to translate key findings to human pathophysiology. The use of high frequency ultrasound technology provided us with clinically relevant parameters to assess changes in maternal, foetal and placental blood flow that can be directly compared to human studies. Here, we have been able to compare physiological data with mechanistic studies of the placenta which is otherwise a major challenge in clinical research.

### Limitations

Whilst rodent models are commonly used in both reproductive and cardiovascular research, we acknowledge that placenta anatomy and development do differ compared to humans. However, once mature the labyrinth zone is functionally analogous to human chorionic villi, as well as undergoing similar vascular remodelling and continuous angiogenesis throughout the pregnancy which makes the mouse a strong model for the current study.

## Conclusion

This foundational study allowed us to gradually increase CBT over a short period of time to examine the consequences of acute heat exposure at mid-gestation. We have established a heating regime which attempted to reflect the physiological response to extreme heat exposure during pregnancy. Our findings indicate that an acute period of heat stress can overwhelm thermoregulatory mechanisms during pregnancy which induces haemodynamic changes, altering placental and foetal blood flow. Consequently, we observed changes in placental architecture, likely disrupting maternal-foetal exchange. As described by the DoHAD theory, this likely increases lifelong risk of metabolic, cardiovascular and neurological conditions. AHE in pregnancy also induced maternal cardiac dysfunction and increased cardiac and renal fibrosis, potentially indicating long term effects of pregnancy complications on cardiovascular health.

Collectively, this study uniquely brings together changes in pathophysiology in response to acute heat exposure at mid gestation, emphasising the urgent need for continued reproductive research in the face of climate change. There is still a lack of data determining the lifelong impact of prenatal heat stress on offspring that should be addressed in future studies.

## Acknowledgments

We acknowledge the support of the Resource Unit Staff and Dundee Imaging Facility at the University of Dundee for their pivotal support in this project. This study utilized the VevoF2 which was awarded to Dr Colin Murdoch through the Medical Research Council World Class Laboratory Equipment Grant MC_PC_MR/X013057/1. We acknowledge Prof Kate Storey’s scientific discussions regarding the model and technical support with histochemistry.

